# Exploration of synergistic action of cell wall-degrading enzymes against *Mycobacterium tuberculosis*

**DOI:** 10.1101/2021.04.05.438542

**Authors:** Loes van Schie, Katlyn Borgers, Gitte Michielsen, Evelyn Plets, Marnik Vuylsteke, Petra Tiels, Nele Festjens, Nico Callewaert

## Abstract

**Background:** The major global health threat tuberculosis is caused by *Mycobacterium tuberculosis* (Mtb). Mtb has a complex cell envelope – a partially covalently linked composite of polysaccharides, peptidoglycan and lipids, including a mycolic acid layer – which conveys pathogenicity but also protects against antibiotics. Given previous successes in treating gram-positive and -negative infections with cell wall degrading enzymes, we investigated such approach for Mtb.

**Objectives:** (i) Development of an Mtb microtiter growth inhibition assay that allows undisturbed cell envelope formation, to overcome the invalidation of results by typical clumped Mtb-growth in surfactant-free assays. (ii) Exploring anti-Mtb potency of cell wall layer-degrading enzymes. (iii) Investigation of the concerted action of several such enzymes.

**Methods:** We inserted a bacterial luciferase-operon in an auxotrophic Mtb strain to develop a microtiter assay that allows proper evaluation of cell wall degrading anti-Mtb enzymes. We assessed growth-inhibition by enzymes (recombinant mycobacteriophage mycolic acid esterase (LysB), fungal α-amylase and human and chicken egg white lysozymes) and combinations thereof, in presence or absence of biopharmaceutically acceptable surfactant.

**Results:** Our biosafety level-2 assay identified both LysB and lysozymes as potent Mtb-inhibitors, but only in presence of surfactant. Moreover, most potent disruption of the mycolic acid hydrophobic barrier was obtained by the highly synergistic combination of LysB, α-amylase and polysorbate 80.

**Conclusions:** Synergistically acting cell wall degrading enzymes are potently inhibiting Mtb – which sets the scene for the design of specifically tailored antimycobacterial (fusion) enzymes. Airway delivery of protein therapeutics has already been established and should be studied in animal models for active TB.

## Introduction

Pathogens of the genus *Mycobacterium* are the etiological agents of various severe diseases, the most notorious being tuberculosis (TB) caused by *Mycobacterium tuberculosis* (Mtb). It is the most lethal infectious disease worldwide and while it might be surpassed by COVID-19 in 2020, it still poses a major global health threat – and worryingly, multidrug-resistance is on the rise. Treatment success of multi-drug resistant tuberculosis (MDR-TB) is currently only 57% and the very protracted treatment comes with harsh side-effects.^1^ Recent advances in drug development have drastically shortened the treatment of MDR-TB and provided more tolerable oral regimens composed of both established TB drugs, novel compounds and repurposed drugs.^1–5^ Still, MDR-TB treatment typically takes 9 to 20 months and the search for more effective and quicker acting drugs is ongoing. Main challenges are the slow growth of Mtb and the relatively impermeable, complex layered cell wall structure of mycobacteria (Figure 1), in which the cell membrane is covered by a layer of peptidoglycan (PG) covalently linked to arabinogalactan. Mycolic acids are esterified to the AG and together with free, intercalating (glyco- and phospho-) lipids and fatty acids, they constitute a near impermeable outer ‘mycomembrane’.^6–9^ This mycomembrane is surrounded by a capsule comprised mainly of α-glucan.^10,11^

**Figure 1.**
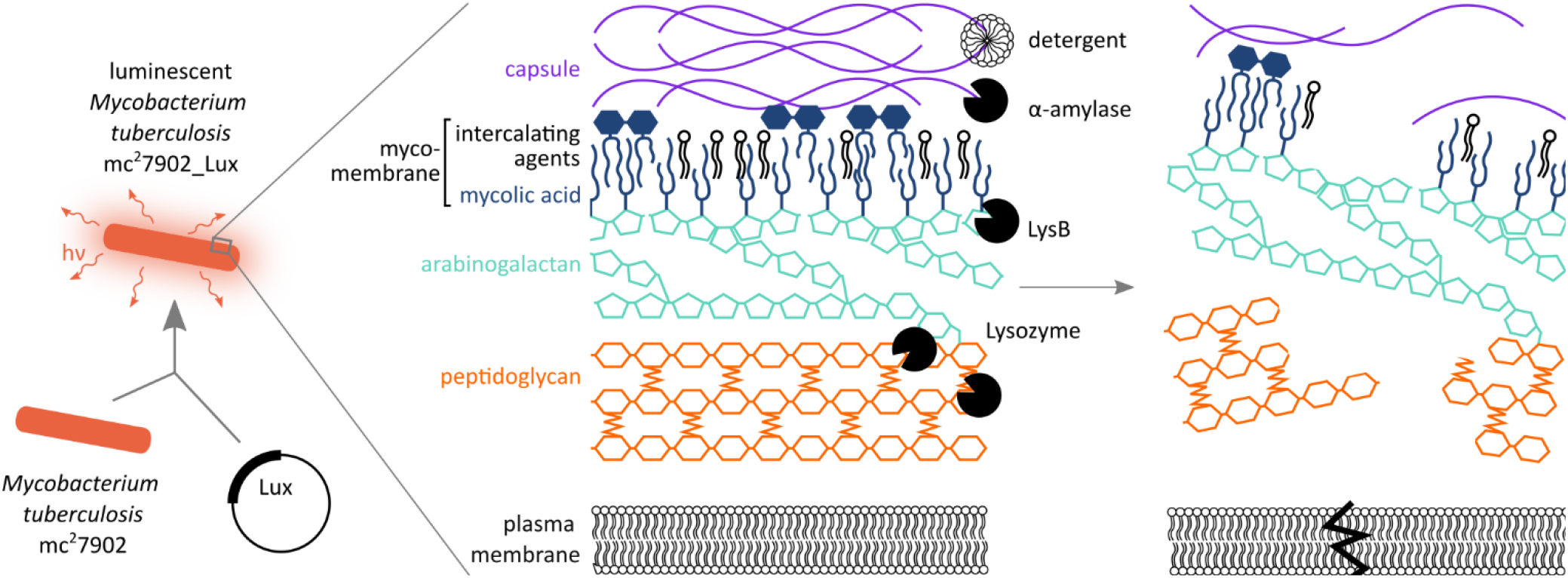
Exploring the synergistic action of cell-wall degrading enzymes to permeabilize the *Mycobacterium tuberculosis* cell wall. Strategy: the triple auxotrophic Mtb mc^2^7902 strain is transformed with a bacterial Lux-operon^**34**^ to allow for the use of a bioluminescent assay to test drug sensitivity. Putative cell wall-degrading agents are added alone or in combination in order to destabilize the structural layers of the cell envelope, leading to permeabilization. Polysorbate 80 is a non-ionic surfactant; α-amylase is an α-glucan-hydrolyzing enzyme; LysB is a mycolylarabinogalactan esterase and lysozyme is a peptidoglycan-hydrolyzing enzyme. Intercalating (glyco- and lipo-)proteins in the cell wall layers are not shown to maintain clarity.

Bacteriophage PG-degrading enzymes (endolysins) are increasingly investigated as antibacterial agents, with several products for topical use against Gram-positive infections already on the market,^12,13^ and progress being made for Gram-negative pathogens.^14–16^ Such products are rapidly gaining attention due to the looming antibiotics resistance crisis, together with improved know-how in biopharmaceutical protein production and formulation for nebulization or dry powder inhalation. In the context of respiratory diseases such as TB, it is encouraging that several recombinant biopharmaceutical protein treatments (e.g. dornase alpha to reduce viscosity of airway mucus) have been successfully developed for inhalation.^17^ However, due to their distinctive cell wall, bacteriophage-mediated lysis of mycobacteria is more complicated than that of other Gram-positives. In addition to endolysin, mycobacteriophages employ a mycomembrane-targeting mycolylarabinogalactan esterase (LysB) to lyse mycobacteria.^18–20^

Whereas endolysin derived from mycobacteriophage Ms6 has been reported not to inhibit *M. smegmatis* or Mtb growth when added therapeutically,^21^ the smaller hen egg white lysozyme (which also cleaves PG) has long been shown to weakly inhibit both *M. smegmatis* and Mtb.^22–25^ For *M. smegmatis*, a growth-inhibitory effect of LysB enzymes has been described in presence of surfactant or membrane-destabilizing cationic peptides.^18–20,26^ Hen egg white lysozyme, RipA (an Mtb PG-endopeptidase), RpfE (an Mtb putative transglycosylase) or hydrolase-30 (an *M. smegmatis* cell wall hydrolase) had moderate inhibitory effects on the growth of *M. smegmatis*, but this effect was significantly increased if the enzymes were administered in combination with various antibiotics, confirming the role of the mycomembrane as a drug barrier.^25^ This illustrates how cell wall-weakening enzyme treatments have the potential to significantly shorten the treatment regimens of Mtb chemotherapy.

Previous studies on the use of cell wall degrading enzymes against Mycobacteria suffer from two main limitations. First, these studies often use *Mycobacterium smegmatis* as the test organism, a rather distant relative to Mtb, with known differences in cell wall composition.^27,28^ This organism is non-infectious and grows very rapidly, hence its popularity. However, for cell wall degrading antibiotics research the pathogen should be used. Second, Mtb clumps heavily in cultures due to the composition of its cell wall. As this has precluded most high-throughput drug test formats, the surfactant polysorbate 80 is routinely added to Mtb culture medium to support finely dispersed planktonic growth. Polysorbate 80 does not restrict Mtb propagation in cultures that use glycerol as the carbon source.^29,30^ Still, it is known to alter mycobacterial drug susceptibility^31,32^ and to destabilize the outer capsule.^11^ Obviously this could affect the validity of results obtained with cell wall degrading enzymes, and precludes exploration of surface-active components in the enzyme cocktail. In this study, we solve both problems by developing a bioluminescent derivative of a biosafety level 2 triple auxotrophic *Mtb*.^33^ The bacterial lux operon used does not require cellular uptake of any cofactor,^34^ which allows for quantifying metabolic activity in static, clumped cultures.

We set out to systematically assess the potency of enzymes that degrade the various layers of the Mtb cell wall, using cultivation conditions that keep the Mtb cell wall/biofilm intact. Apart from PG and the mycomembrane, we targeted the outer capsule of *Mtb*. Considering that α-glucan is a main constituent of this capsule,^10^ we speculated that *Aspergillus oryzae* α-amylase, an α-glucan hydrolyzing enzyme already used in human medicine to treat pancreatic insufficiency, could have a capsule-destabilizing effect and potentially synergise with the known capsule destabilization imparted by polysorbate 80. Furthermore, we hypothesized that the anti-TB potency of cell wall-degrading enzyme treatments would be enhanced by using the enzymes in cocktails, which could potentially ‘peel’ the cell envelope layer-by-layer and synergistically weaken it (Figure 1). Amylase-induced hydrolysis of the capsular layer may, for instance, increase permeability for enzymes such as LysB that degrade the mycomembrane and/or for enzymes such as lysozyme that degrade the PG layer underneath.

## Materials and Methods

### Materials

Chicken egg white lysozyme was purchased at Sigma-Aldrich (L6876, ≥90% protein), as was human lysozyme recombinant from rice (L1667, ≥90% pure) and *Aspergillus oryzae* α-amylase (A8220, ≥800 fungal amylase units/g). All were dissolved in phosphate-buffered saline (PBS) and protein concentrations were determined via spectrophotometry at 280 nm. Polysorbate 80 (Sigma-Aldrich) was dissolved at 0.2% in PBS. Antibiotics isoniazid (INH) and rifampicin (RIF) (Sigma-Aldrich) were dissolved as stock solutions of 1 mg/ml (INH) in PBS and 10 mg/ml (RIF) in methanol. All solutions were 0.22 µm filter-sterilized.

### Plasmid construction and enzyme expression

Sequences encoding LysB originating from mycobacteriophages Ms6 and D29 (LysB-Ms6 and LysB-D29, UniProt accession numbers Q9FZR9 and O64205, respectively) were ordered as synthetic DNA in the pUC57 vector (GenScript). Via PCR, the genes were cloned into the pLH36 vector in frame with an N-terminal His6-tag flanked by a murine caspase-3 cleavage site (sequences in Supplementary A). The expression constructs were verified by Sanger sequencing (VIB Genetic Service Facility, Antwerp, Belgium) and transformed to *E. coli* BL21 (DE3). After initial growth at 28 °C in terrific broth medium (Sigma-Aldrich) supplemented with carbenicillin, expression was induced at a culture OD_600_ of 0.6 by the addition of 0.6 mM isopropyl-β-D-thiogalactopyranoside. LysB-Ms6 was expressed overnight at 28 °C and LysB-D29 for 4h at 37°C. Bacteria were harvested by centrifugation (8000x*g*, 15 min at 4 °C).

### Purification of LysB enzymes

After resuspension, bacterial pellets were incubated for 30 min in ice-cold lysis buffer (25 mM Tris-HCl pH 8, 5 mM MgCl_2_, 0.1% Triton X-100 and 1X cOmplete™ protease inhibitor (Roche)) and lysed by sonication at 70% amplitude using a Qsonica sonicator (4s on, 8s off for 9 min). After 30 min centrifugation (100,000x*g*) at 4 °C, cleared lysates were 0.22 µm filter sterilized. Enzymes were isolated from the cleared lysates via nickel affinity chromatography (IMAC) using a HisTrap column (GE Healthcare) and size exclusion chromatography (SEC) using a SuperDex 75 10/300 or 16/600 SEC column (GE Healthcare) calibrated with 20 mM HEPES pH 7 and 150 mM NaCl. After quantification (BCA assay, Pierce), enzymes were 0.22 µm filter sterilized and aliquots were snap-frozen in liquid nitrogen before storage at -80 °C.

### Evaluation of enzyme activity

Lipolytic LysB activity was quantified using an assay adapted from literature.^18,35^ Briefly, LysB was incubated with 5 mM *p*-nitrophenol butyrate substrate (pNPB) in 20 mM Tris-HCl pH 8.0, 100 mM NaCl and 0.1% Triton X-100. The release of *p*-nitrophenol after hydrolysis of the pNPB substrate was measured as increase in absorbance at 400 nm for 30 minutes, and the ΔA_400nm_/minute was calculated from the linear region of the blank-corrected values after a lag-phase. The specific enzyme activity was calculated using the micromolar extinction coefficient of *p*-nitrophenol at 400 nm at 37°C of 0.0148. One LysB enzyme unit releases 1 nmol of *p*-nitrophenol per minute at pH 8 at 37 °C using pNPB as substrate.

PG-degrading activity of human and hen egg white lysozyme was determined on a fluorescein-labelled *Micrococcus lysodeiticus* cell wall substrate using the EnzChek Lysozyme Assay Kit (ThermoFisher), in which one unit is defined as the amount of enzyme required to produce a 0.001 units per minute change in the absorbance at 450 nm of at pH 6.24 and 25°C, using a suspension of *Micrococcus lysodeikticus* as the substrate.

### Mycobacterium strain and handling

To enable a bioluminescence-based *Mycobacterium* drug susceptibility assay, we used the pMV306hsp+LuxG13 plasmid (a gift from Brian Robertson and Siouxsie Wiles, Addgene #26161) which contains a bacterial luciferase operon enhanced by using the G13 promotor in front of luxC.^36^ The integrase-free reporter plasmid pMV306DIhsp+LuxG13 (BCCM/GeneCorner accession number LMBP11308) was produced by deleting the int gene from pMV306hsp+LuxG13 via inverted PCR using phosphorylated primers (5’-Pi-GTCCATCTTGTTGTCGTAGGTCTG-3’ and 5’-Pi-TCTTGTCAGTACGCGAAGAACCAC-3’),^37^ followed by ligation of the product. The use of an integrase-free reporter plasmid and a separate suicide vector containing integrase (pBlueScript-Integrase, a gift from Peter Sander, Institute of Medical Microbiology, University of Zurich) allowed for temporary integrase activity – to integrate the reporter plasmid – and subsequent selection of integrase-free reporter strains.^38^ The biosafety level 2-approved triple auxotrophic ΔpanCD ΔleuCD ΔargB Mtb mc^2^7902 strain^33^ (a gift from W.R. Jacobs Jr.) was co-transformed with pMV306DIhsp+LuxG13 and pBlueScript-Integrase via electroporation (12.5 kV/cm, 25 μF capacitance, 800 Ω resistance). After selection on kanamycin, the obtained Mtb mc^2^7902_Lux strain was stored in 1 ml aliquots at OD_600_ 1 in 20% glycerol at -80 °C. For each drug testing assay, two Mtb mc^2^7902_Lux aliquots were thawed, combined and cultured at 37 °C, shaking, in a 60 ml square bottle in 20 ml of Middlebrook 7H9 broth (BD Diagnostics) supplemented with 0.05%_V/V_ polysorbate 80, 10% oleic acid-albumin-dextrose-catalase supplement (OADC; BD Diagnostics), 0.5%_V/V_ glycerol, 1 mM L-arginine, 50 µg/ml L-leucine and 24 µg/ml L-pantothenate. At days 5 and 8 after thawing, bacteria were subcultured by 1/100 and 1/20 dilution, respectively, for use in setting up the antimicrobial assays on day 9. Middlebrook 7H10 agar (BD Diagnostics) supplemented with 10% OADC, 0.5%_V/V_ glycerol, 1 mM L-arginine, 50 µg/ml L-leucine and 24 µg/ml L-pantothenate was used for growth of *Mtb* on solid culture; colony counts after incubation at 37 °C for up to 12 weeks were given as colony forming units per ml plated (cfu/ml).

### A bioluminescence microtiter assay for mycobacterial growth inhibition

To remove bacterial clumps before setting up the microtiter assay, mycobacteria were passed three times through a 27G needle. A Mtb mc^2^7902_Lux inoculum at an OD_600_ of 0.008 (2·10^6^ cfu/ml) was prepared in assay medium (Middlebrook 7H9 broth supplemented with 10% OADC, 0.5%_V/V_ glycerol, 1 mM L-arginine, 50 µg/ml L-leucine and 24 µg/ml L-pantothenate) without any polysorbate 80 unless specifically stated otherwise.

Drug susceptibility was tested in a microtiter plate format based on the bioluminescence assay proposed by Andreu *et al* (2012).^34^ Briefly, duplicate twofold serial dilutions of various compounds were prepared in assay medium in sterile 96-well black opaque plates (CulturPlate-96 Black, PerkinElmer). To allow for synergy testing, horizontal twofold dilution series of one compound were combined with vertical twofold dilution series of another compound in a ‘checkerboard’ fashion (Supplementary B). Afterwards, the Mtb mc^2^7902_Lux inoculum was added to 2·10^5^ cfu per well. Each plate contained three wells containing inoculum without drugs (positive growth controls), and three wells containing no inoculum (negative controls). All perimeter wells were filled with 200 µl ultrapure water to limit evaporation from sample wells. Assay plates were incubated at 37 °C in an incubator providing 5% CO_2_and 80% humidity, which yielded a higher reproducibility than when plates were placed (in a sealed box or plastic bag) in a regular incubator (data not shown). Bioluminescence was measured after 1, 4, 7, 10 and 13 days of incubation in a GloMax^®^ 96 or GloMax Navigator Microplate Luminometer (Promega) with 1.0 s integration per well, and expressed as relative light units (RLU).

### Bioluminescence microtiter assay data processing (details on models and synergy evaluation in Supplementary C)

After averaging bioluminescence data of technical duplicate microtiter plates, data of three independent experiments were subjected to fitting of the generalized logistic model and the Gompertz model^39,40^ using GenStat 19.1 (VSN International Ltd). The minimal inhibitory concentration (MIC), defined as the intersection of the lower asymptote with the tangent to the inflexion point of the curve (Figure 3a), was calculated for the best fitting dose-response curve of each dataset (judged by the percentage of variance accounted for by the fitted curve). If no inhibition was observed, the MIC was set as larger than or equal to the highest concentration tested. If inhibition was observed but no dose-response curve could be fitted, the MIC was set as lower than or equal to the lowest value for which all data points were below 200 RLU (corresponding to the bioluminescence of half the inoculum).

For synergy assessment, data were normalized to the means of positive and negative growth controls and evaluated using the Combenefit tool.^41^ Subsequently, a linear mixed model was fitted to further assess the significance of the synergy detected as a statistical interaction between compounds.

Correlation between cfu/well and RLU/well was assessed using the SciPy Stats package.^42,43^

## Results

### Enzyme production and characterization

Mycobacteriophage Ms6 and D29 mycolic acid esterases (LysB-Ms6 and LysB-D29) were produced in *E. coli* and purified via IMAC and SEC to yields of 3.12 and 1.34 mg/liter *E. coli* culture (i.e. 81 and 44 nmol/l), respectively (Figure 2a-b). Minor low molecular weight bands were observed in purified Lys-Ms6, probably indicating low level protein degradation. Enzymatic activity on a pNPB substrate seemed highly dependent on the bacteriophage from which the enzyme was derived, with LysB-D29 displaying three-fold higher specific activity than LysB-Ms6 (Table 1, not significant, p=0.2, Mann-Whitney test). Enzymatic activity of commercially available lysozyme was validated using a *Micrococcus* PG degradation assay (Table 1, bottom).

**Table 1.** *In vitro* activity-testing of potential antimycobacterial enzymes. Lipolytic activity of LysB enzymes was determined in vitro on a pNP-butyrate substrate. One LysB enzyme unit (U) will release 1 nmol of p-nitrophenol per minute at pH 8 at 37 °C. Peptidoglycan (PG)-degrading activity of human and hen egg white lysozyme was determined on a fluorescein-labeled Micrococcus lysodeiticus cell wall substrate. One lysozyme enzyme unit (U) will produce a 0.001 units per minute change in the absorbance at 450 nm of at pH 6.24 and 25°C.

| Compound | Substrate | *In vitro* substrate-degrading activity |  |
| --- | --- | --- | --- |
|  |  | U/nmol | n |
| LysB-Ms6 | pNP-butyrate | 338 ± 7.7 | 3 |
| LysB-D29 |  | 1375 ± 173.7 | 2 |
| Lysozyme (hen egg white) | *Micrococcus* peptidoglycan | 771 ± 26.1 | 3 |
| Lysozyme (human) |  | 589 ± 13.2 | 3 |

**Figure 2.**
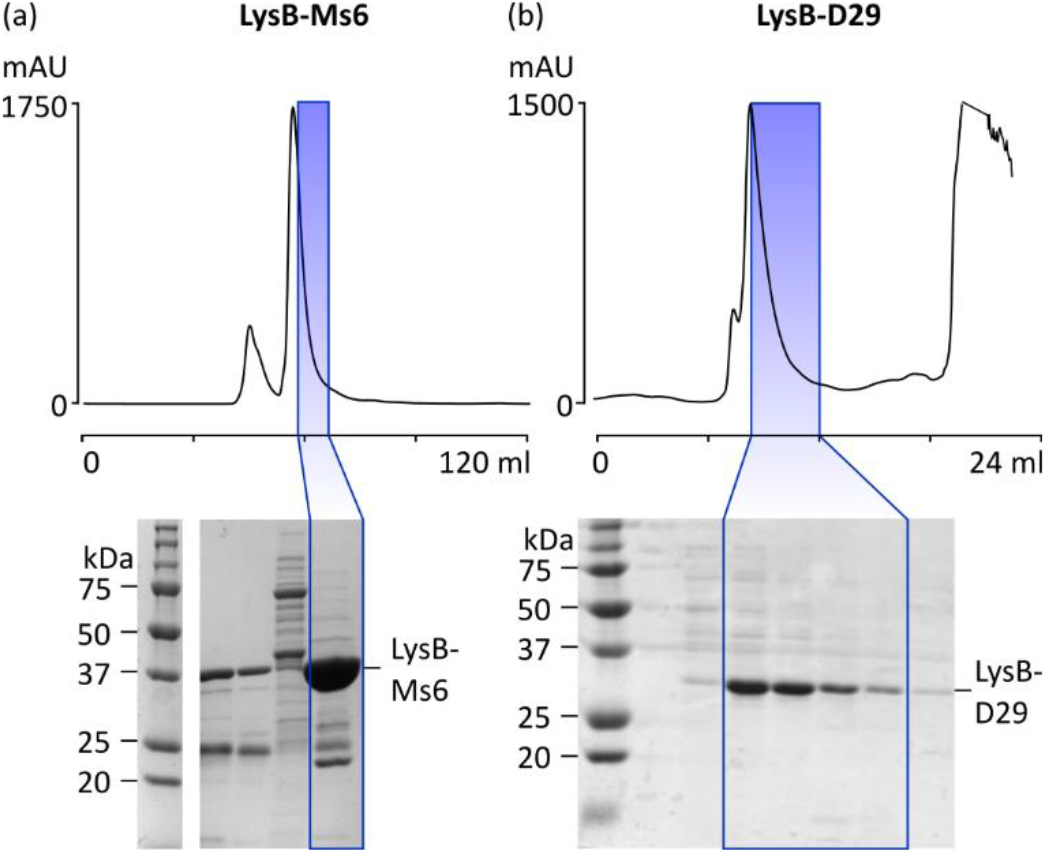
Production of (a) LysB-Ms6 and (b) LysB-D29. After expression in E. coli BL21 (DE3), proteins were purified via nickel chromatography and size exclusion chromatography (SEC, chromatogram shown). After SEC, peak fractions containing the recombinant protein at highest purity were pooled, indicated in this figure by blue rectangles on SEC chromatograms and Coomassie-stained SDS-PAGE.

### Cell wall degrading enzymes inhibit mycobacterial growth

Lux operon-induced bioluminescence has already been shown to correlate very well with (inhibition of) mycobacterial growth in drug susceptibility assays.^34,36^ We therefore used this bioluminescence to reliably evaluate the antimycobacterial activity of recombinant LysB-Ms6 and LysB-D29 as well as commercially available lysozyme (human and chicken egg white) and α-amylase (*A. oryzae*) and antibiotics isoniazid (INH) and rifampicin (RIF). Bioluminescence was determined after 1, 4, 7, 10 and 14 days of incubation. When modeling dose-response curves, inhibition often appeared incomplete at day 1 (parameter A not zero) and after excluding that time point, the highest percentage of explained variance was consistently obtained at day 4. Therefore, minimal inhibitory concentrations (MICs) were determined at this time point (Table 2 and Figure 3b).

**Table 2.** MIC values of compounds against Mtb mc^2^7902_Lux. Luminescence drug susceptibility assay (n = 3 with technical duplicates). MIC was determined according to Lambert & Pearson (2000)^**39**^. If no inhibition was observed, the MIC was defined as larger than or equal to the highest concentration tested. If inhibition was observed but no sigmoid could be fitted, the MIC was defined as lower than or equal to the lowest value for which all datapoints were lower than 200 RLU (corresponding to half the inoculum).

| Compound | MIC (μM) and standard error (s.e.) |  | MIC fold-decrease |
| --- | --- | --- | --- |
|  | in assay medium | in assay medium + 0.05% polysorbate 80 |  |
| Rifampicin | 0.3 ± 0.06 | ≤ 0.02 | ≥ 10 |
| Isoniazid | 0.1 ± 0.03 | 0.2 ± 0.03 | 0.6 |
| α-Amylase (*A. oryzae*) | ≥ 250 | ≥ 250 | N/A |
| Lysozyme (hen egg white) | ≥ 250 | 40 ± 5 | ≥ 6 |
| Lysozyme (human) | ≥ 250 | 33 ± 4 | ≥ 8 |
| LysB-D29 | ≥ 0.4 | 0.08 ± 0.01 | ≥ 5 |
| LysB-Ms6 | ≥ 0.4 | 0.2 ± 0.07 | ≥ 2 |

We calculated a MIC of INH in the same order of magnitude as described in Andreu *et al* at day 4.^34^ The MIC we calculated for RIF was approximately ten-fold higher, but it should be noted we used a different strain of Mtb (pathogenic H37Rv in literature versus mc^2^7902 in our study) and our definition of MIC is more stringent than that used in literature (1 log reduction in bioluminescence).

Both human and chicken egg white lysozyme were only marginally inhibiting mycobacterial growth by themselves, but become equally effective (p=0.318; t-test) in presence of polysorbate 80, with activities enhanced 6 to 8-fold (Figure 3b and Table 2). α-Amylase, on the other hand, did not considerably affect growth in the concentration range tested, neither in the presence nor in the absence of surfactant (MIC ≥250 µM). LysB-D29 and LysB-Ms6 could inhibit growth equally effectively in presence of polysorbate 80 (MIC= 0.08 ± 0.01 µM and 0.2 ± 0.07 µM, respectively; t-test p=0.189). It should be noted that the dose-response curve shapes differ, with LysB-D29 showing a sharp dose-dependent effect and stronger polysorbate 80-dependence than LysB-Ms6.

**Figure 3.**
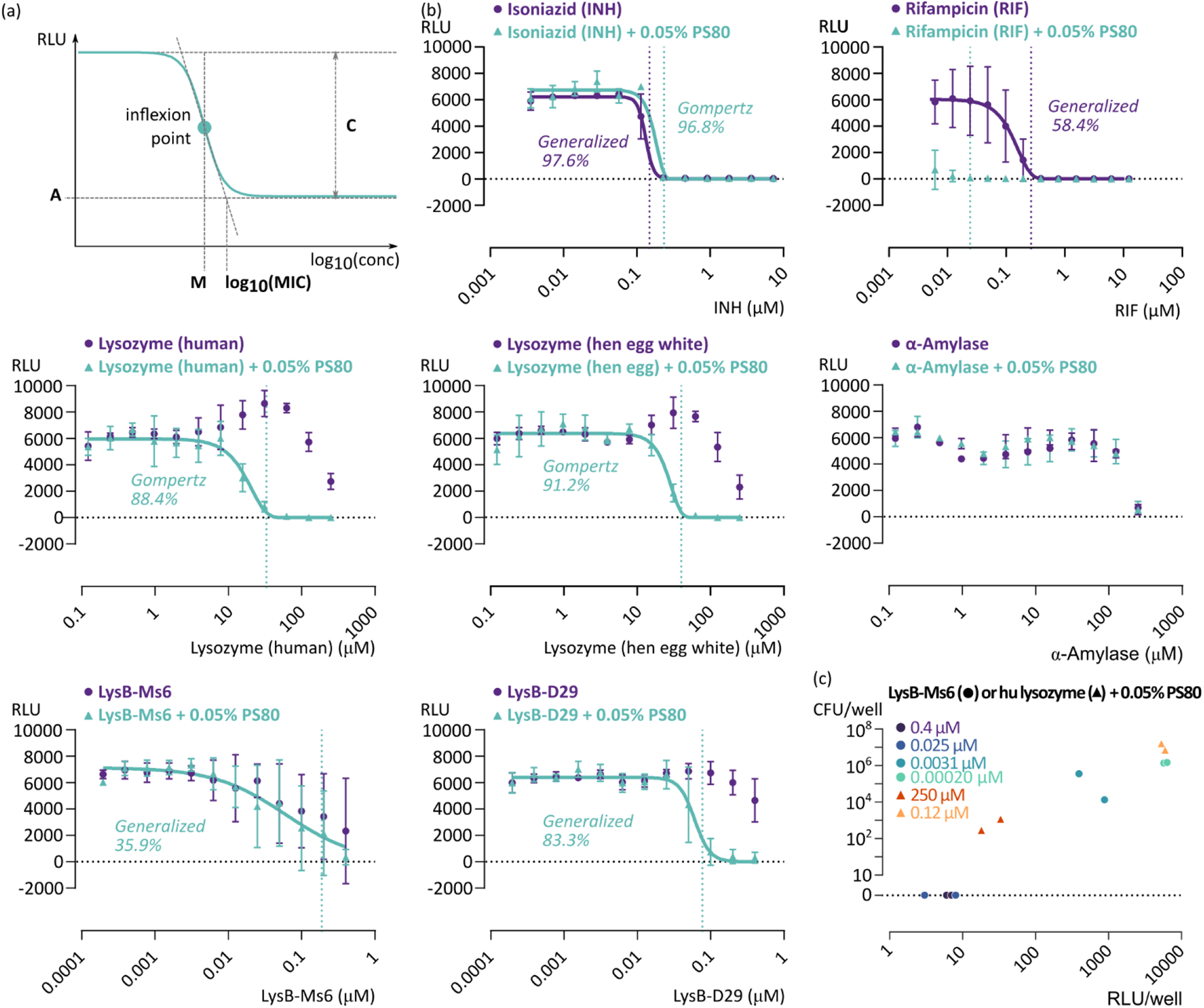
The effect of various antimycobacterial compounds on bioluminescence of Mtb mc^2^7902_Lux. **(a)** Scheme of the generalized logistic function as applied to a bioluminescence growth assay and its main parameters. **(b)** Effect of antimycobacterial agents in absence (purple circles) and presence (cyan triangles) of 0.05% polysorbate 80 (PS80). Luminescence detected after 4 days of incubation at 37 °C. If data allowed, the generalized logistic model and the Gompertz model were fitted to the luminescence data (the model explaning the highest percentage of variance is shown). The MIC calculated from regression parameters according to Lambert & Pearson (2000)^**39**^ is indicated as a dotted vertical line. If no inhibition was observed, the MIC was defined as larger than or equal to the highest concentration tested (not shown in graph). If inhibition was observed but no sigmoid curve could be validly fitted, the MIC was defined as lower than or equal to the lowest value for which all datapoints were lower than 200 RLU (corresponding to half the inoculum). Data shown were derived from three independent experiments, with each data point an average of duplicate plates within a repetition of the experiment. Mean and standard deviation are shown. **(c)** For several conditions from the high and low ends of inhibition curves of LysB-Ms6 (circles) and human lysozyme (triangles), both in presence of 0.05% PS80, viability was determined by cfu plating on solid medium after measuring luminescence. Datapoints shown are averages of technical plating replicates.

To assess viability of growth-inhibited mycobacteria, several conditions at the high and low ends of the LysB-Ms6 and human lysozyme inhibition curves were selected for a repeat luminescence assay followed by cfu determination, which demonstrated a strong correlation between RLU and cfu (Figure 3c, Spearman correlation coefficient = 0.93, p=1.6·10^−5^). Upon administration of 250 µM human lysozyme in presence of 0.05% PS80, viability was vigorously decreased after 4 days at 37 °C (10^3^ cfu/well versus 10^7^ cfu/well after sub-MIC treatment) and upon administration of the highest concentrations of LysB-Ms6 with 0.05% PS80, no viable *Mtb* were observed at all (Figure 3c).

### Synergy

Considering the layered structure of the mycobacterial cell wall, we hypothesized that cell wall degrading enzymes can display synergy if administered jointly. Using Combenefit^41^ we analysed various combinations of antimycobacterial compounds in checkerboard assays (Table 3). The three tested compound combinations scoring highest for synergy were selected for in-depth analysis: LysB-Ms6 with human lysozyme and 0.05% polysorbate 80, and LysB-Ms6 with α-amylase in presence or absence of 0.05% polysorbate 80. When LysB-Ms6 was combined with either human lysozyme or *A. oryzae* α-amylase, both synergistic and antagonistic regions were observed in the dose-response surface with few significant combinations and an overall neutral effect (Figure 4a-b). However, when 0.05% polysorbate 80 was added to the combination of LysB-Ms6 and *A. oryzae* α-amylase, the shape of the dose-response surface changed drastically, displaying a deep valley of synergy in which 18 out of 36 conditions scored significant for synergy (p<0.05 to p<0.0001; Figure 4c).

**Table 3.** Synergy and antagonism of antimycobacterial compounds as determined by the Combenefit^41^ tool. Luminescence drug susceptibility assay on Mtb mc^2^7902_Lux (n = 3 with technical duplicates). Compounds were added ‘checkerboard’-wise in combined dilution series, of which the highest concentration is indicated. The half maximal inhibitory concentration (IC50) was derived from a Hill equation fit of the data. Synergy and antagonism scores were determined using either the Loewe, Bliss or HSA model. Shading indicates three highest scoring synergy conditions. Lysozyme: human; α-amylase: *Aspergillus oryzae*. If present as compound C, polysorbate 80 is at 0.05%.

| Compound |  |  | Max conc (μM) |  | IC<sub>50</sub> (μM) |  | Sum antagonism score |  |  | Sum synergy score |  |  |
| --- | --- | --- | --- | --- | --- | --- | --- | --- | --- | --- | --- | --- |
| A | B | C | A | B | A | B | Loewe | Bliss | HSA | Loewe | Bliss | HSA |
| α-Amylase | LysB-Ms6 | - | 125 | 0.2 | 125.00 | 0.13 | -11.51 | -10.63 | -10.37 | 27.73 | 33.09 | 35.10 |
| α-Amylase | LysB-Ms6 | polysorbate 80 | 125 | 0.2 | 0.04 | 0.05 | 0.00 | 0.00 | 0.00 | 117.06 | 100.37 | 117.31 |
| Lysozyme | LysB-D29 | polysorbate 80 | 125 | 0.2 | 14.90 | 0.04 | -34.14 | -24.17 | -18.05 | 0.00 | 1.34 | 2.93 |
| Lysozyme | LysB-Ms6 | - | 125 | 0.2 | 125.00 | 0.04 | -49.85 | -48.63 | -48.63 | 7.93 | 8.71 | 8.95 |
| Lysozyme | LysB-Ms6 | polysorbate 80 | 125 | 0.2 | 18.75 | 0.14 | -17.71 | -17.88 | -17.63 | 36.78 | 46.89 | 53.27 |

**Figure 4.**
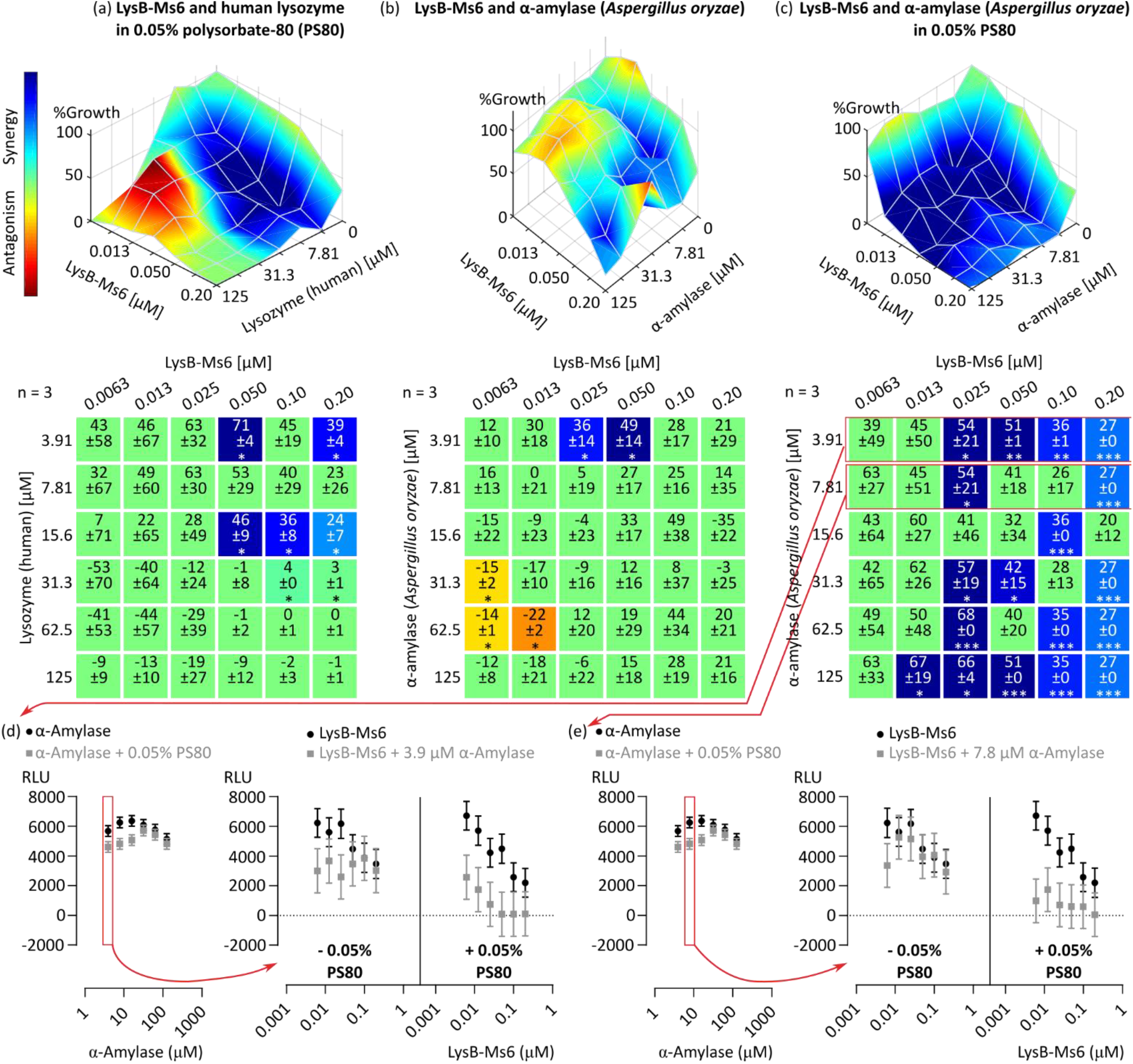
(a, b and c) Synergy of various antimycobacterial compounds (Bliss model) mapped to the experimental dose-response curve, and the corresponding synergy and antagonism matrix. Data were derived from a checkerboard bioluminescence drug susceptibility assay of Mtb mc^2^7902_Lux. Bioluminescence was detected after 4 days of incubation at 37 °C. Data shown were derived from 3 independent experiments, each with duplicate technical replicate plates. The synergy score shown in the matrix is the percentage of growth inhibition by the drug combination that is not explained by the Bliss-modeled reference dose-response, as calculated using the Combenefit^**41**^ tool. Results are coloured according to the obtained synergy score only if the result is significant following a one-sample t-test. *p<0.05 **p<0.001 ***p<0.0001. **(d and e) Mixed model of the synergy of LysB-Ms6 with α-amylase (*A. oryzae*) in the presence or absence of 0**.**05% polysorbate 80**. A linear mixed model was applied to the bioluminescence data of two sets of drug combinations from (c), to evaluate the separate and combined effects of each component (LysB-Ms6, α-amylase or PS80) on the variance. For each set of drug combinations, the resulted predicted values are plotted: the effect of α-amylase alone or in presence of PS80 (left) and the combined effect of LysB-Ms6 and a set concentration of α-amylase in absence (middle) or presence (right) of PS80.

The significance of synergy between LysB-Ms6, α-amylase and polysorbate 80 was further assessed by applying a linear mixed model to bioluminescence data of two sets of drug combinations (Figure 4d-e, details in Supplementary E). In presence of 0.05% polysorbate 80, a significant interaction (p=0.012 in the mixed model) was identified between a dilution series of LysB-Ms6 and 7.8 µM α-amylase, confirming synergy (Figure 4e). In absence of polysorbate 80, no such significant interaction was observed. In the presence of 3.9 µM α-amylase (Figure 4d) a similar trend was observed but it was of borderline significance (p=0.058 in the mixed model). From combined results of bioluminescence assays and synergy analyses, we conclude that the combination of dilution series of LysB-Ms6 and *A. oryzae* α-amylase in the presence of 0.05% polysorbate 80 leads to a strong, synergistic effect on the growth of Mtb mc^2^7902, validating the hypothesis that targeting multiple layers of the cell wall can lead to more effective antimycobacterial activity.

## Discussion

The results of this study can be summarized as a methodological improvement in anti-Mtb drug screening, and a main discovery with regard to the synergistic antimycobacterial effect of cell wall-degrading enzymes.

The methodological improvement pertains to the rather mundane, but vexing problem that Mtb grows in clumps when its cell envelope is left unperturbed. While adding surfactants in high-throughput microtiterplate-based drug studies enables spectrophotometry-based readouts of growth, it likely invalidates susceptibility tests of cell wall-targeting enzymes and drugs, as accessibility to the cell wall substrates of such enzymes will be altered – which we confirmed in our studies. Other researchers have deferred to the use of model organism *M. smegmatis*, which clumps less, grows more rapidly than *Mtb* and is non-infectious, allowing work outside highly restricted and expensive biosafety level 3 laboratories. Nevertheless, the cell wall of *M. smegmatis* differs significantly from that of the pathogenic slow-growing mycobacteria and it is questionable whether results can be extrapolated to Mtb. For this reason, we resorted to the use of Mtb engineered with a bacterial lux operon. In a parallel project, other luciferase systems were also implemented, and whereas higher specific luminescence could be obtained with firefly luciferase, the bacterial lux operon does not require addition of the luciferin cofactor. This is a major advantage, as it eliminates doubts about false results caused by bacterial clumping limiting cofactor diffusion, or cell wall degrading enzyme treatment ‘opening up’ access for such cofactor. Moreover, we combined this system with biosafety convenience by implementing it in a triple auxotrophic Mtb derivative generated in and generously provided by the Jacobs lab.^33^ The luminescent auxotrophic Mtb strain and associated protocols should be useful in a great variety of drug screening studies.

Using this strain, we discovered that the combination of LysB-Ms6 and *A. oryzae* α-amylase was highly synergistic in inhibiting mycobacterial growth in presence of polysorbate 80. Remarkably, α-amylase itself has no anti-Mtb effect either with or without 0.05% polysorbate 80. The most straightforward explanation is that polysorbate 80 and α-amylase collaborate in disrupting the α-glucan capsule (non-essential for *in vitro Mtb* growth),^11^ clearing the way for LysB to hydrolyze the linkage between underlying mycolic acids and the arabinogalactan layer, thus disrupting the mycomembrane. This finding lays the first stone for the creation of specifically tailored antimycobacterial (fusion) enzymes.

The anti-Mtb potential of LysB-Ms6 itself and its improvement in the presence of polysorbate 80 had already been shown on the *M. smegmatis* model, and it was demonstrated that cell wall hydrolysis was the main mechanism behind inhibitory activity of LysB-Ms6.^31^ We report a similar surfactant-dependent bacteriostatic effect of both LysB-Ms6 and LysB-D29 on Mtb.

The mycobacterial cell envelope is a complex non-proteinaceous (thus not directly genetically encoded) structure synthesized by an extensive enzyme machinery.^7^ This likely substantially reduces the chance that single spontaneous mutations lead to resistance against enzyme antimicrobials, contrary to small molecule antibiotics, which bind to genetically encoded protein targets. In addition, it has been observed that in treating MDR-TB patients (the main target group for biopharmaceutical enzyme treatment), low-level antibiotics resistance can be overcome by administrating drug doses exceeding the MICs of the resistant strains.^44^ After weakening the *Mycobacterium* envelope with cell-wall degrading enzymes, classic antibiotics could potentially penetrate more readily, increasing the effective drug concentration intracytoplasmatically to levels above the MIC.

While promising as antimycobacterials, adequate delivery of the enzyme therapeutics to the site of infection is essential to therapeutic efficacy. We believe our study is timely as several inhaled nebulized enzyme therapeutics are under clinical evaluation, providing evidence that lung delivery of therapeutic enzymes is feasible.^17,45^ As an example, it was calculated that 8.8±1.9 mg of an antibody could be delivered to the alveoli of non-human primates using a novel device for protein inhalation.^46^ Considering a conservative estimate of 70 ml lung lining fluid volume,^47^ this would amount to a dose of 0.13 mg/ml, or 4.3 µM of a 30-kDa protein, widely surpassing the MICs we found for LysB in presence of surfactant (polysorbate 80, which has been successfully included in inhaled therapeutics).^17,48^ It should however be considered that mucus hypersecretion is a common feature of airway inflammation, possibly lowering the effective therapeutic dose when treating (MDR-)TB patients.^49,50^ While optimizing lung delivery, topical application of antimycobacterial enzyme therapy might already suffice to treat the skin infection Buruli ulcer caused by *Mycobacterium ulcerans*.^51^

The repeated inhalation of non-human proteins might lead to an immune response, although tolerance is the default response to antigens that come into contact with the mucosa in the deep airways.^52^ If necessary, enzymes could be administered in controlled-release formulations to reduce the number of administration required. Novel drug regimens have already drastically shortened (MDR-)TB therapy but still, duration, complexity and toxicity of (second-line) drug regimens are main challenges in the worldwide fight to end TB.^1^ Enhancing or accelerating the effectiveness of conventional antibiotics by adding cell wall-degrading enzymes in two to three rounds of (controlled-release) enzyme administration, could potentially lower the treatment duration critically and help alleviate the immense burden of TB therapy.

## Acknowledgements

We thank the W.R. Jacobs lab for kindly providing the Mtb mc^2^7902 strain and Dr. B. Robertson, Dr. S. Wiles and Dr P. Sander for expression plasmids used in this study.

## Transparency declarations

None to declare.

